# Effects of healthy aging on tongue-jaw kinematics during feeding behavior in rhesus macaques

**DOI:** 10.1101/2024.07.31.605680

**Authors:** Shreyas Punacha, Kevin Huang, Fritzie I. Arce-McShane

**Author notes:** **Corresponding author:** Fritzie I. Arce-McShane, Department of Oral Health Sciences, School of Dentistry, University of Washington, Seattle, WA, USA.

## Abstract

Several age-related oral health problems have been associated with neurodegenerative diseases such as Alzheimer’s Disease (AD), yet how oromotor dysfunction in healthy aging differ from those found in pathological aging is still unknown. This is partly because changes in the cortical and biomechanical (“neuromechanical”) control of oromotor behavior in healthy aging are poorly understood. To this end, we investigated the natural feeding behavior of young and aged rhesus macaques (Macaca mulatta) to understand the age-related differences in tongue and jaw kinematics. We tracked tongue and jaw movements in 3*D* using high-resolution biplanar videoradiography and X-ray Reconstruction of Moving Morphology (XROMM). Older subjects exhibited a reduced stereotypy in tongue movements during chews and a greater lag in tongue movements relative to jaw movements compared to younger subjects. Overall, our findings reveal age-related changes in tongue and jaw kinematics, which may indicate impaired tongue-jaw coordination. Our results have important implications for the discovery of potential neuromechanical biomarkers for early diagnosis of AD.

## Introduction

Human feeding relies on the coordination of tongue and jaw movements and the precise control of the generation of tongue and bite forces during chews and swallows. Dysfunctions in chewing and swallowing are prevalent in neurodegenerative diseases such as Parkinson’s Disease (PD) and Alzheimer’s Disease (AD), with incidence and severity increasing with age (Dominy et al. 2019; Paganini-Hill et al. 2012; Rogus-Pulia et al. 2015; Seçil et al. 2016; Watanabe et al. 2015). However, age-related effects on the sensorimotor control of feeding behavior is still largely unknown because of a fundamental knowledge gap on the neuromechanical principles that underlie this complex and vital function.

Mastication is a sensorimotor activity essential for preparing food for swallowing. It consists of rhythmically occurring gape cycles, with each cycle consisting of a jaw-opening phase followed by a jaw-closing phase. Although mastication has a typical form and rhythm, it is nevertheless adapted to various properties of food, such as hardness (Hiiemae and Palmer 1999), bite size (Hiiemae et al. 1996), and texture (Schindler et al. 1998), for efficient food breakdown and digestion. The rhythmic activity in chewing is orchestrated by the masticatory central pattern generator (CPG), a network of neurons located in the brainstem (Kohyama et al. 2002; Lund and Kolta 2006; Mistry and Hamdy 2008; Morquette et al. 2012; Nakamura and Katakura 1995). The sensorimotor cortex is also involved in planning and executing precise motor actions, providing the necessary control for dynamic adjustments required during chewing and swallowing. The basal ganglia contributes to the regulation of movement initiation and amplitude, while the cerebellum integrates sensory feedback to fine-tune motor output. Altogether, these brain regions work together towards adaptability and accuracy of masticatory actions in response to varying food properties and external stimuli (Avivi-Arber and Sessle 2018; Lund and Kolta 2006; Morquette et al. 2012; Sessle 2006).

In healthy aging, there is a higher incidence of tooth loss (Ueno et al. 2008), decreased bite force (Bakke et al. 1990; Hatch et al. 2001; Helkimo et al. 1977), loss of jaw muscle area and density (Newton et al. 1993), and weakened orofacial muscles (Newton 1987). Behavioral changes, such as decline in tongue motor skills (Koshino et al. 1997) and masticatory efficiency (Kohyama et al. 2002; Österberg et al. 1996; Wayler and Chauncey 1983), increase with age (Peyron et al. 2004). These age-related changes impact bolus formation, chewing, swallowing (Logemann 1990; Prinz and Lucas 1997; Daniels et al. 2004), and the ability to clear residual debris in the mouth and throat (Yoshikawa et al. 2005). Sensorimotor changes in the periphery are accompanied by age-related changes in brain activity (Lenz et al. 2012; Sebastián and Ballesteros 2012; Brodoehl et al. 2013; Kalisch et al. 2009; Talelli et al. 2008; Bhandari et al. 2016; Ward and Frackowiak 2003; Heuninckx et al.2008; Hutchinson et al. 2002; Seidler et al. 2015; Cheng and Lin 2013; Heise et al. 2013; Cheng et al. 2015; Rueda-Delgado et al. 2019; Fujiyama et al. 2012; McGinley et al. 2010). From a sensorimotor control perspective, these may impact the ability of neurons in the orofacial sensorimotor cortex to control coordination of the tongue and jaw movements (Arce-McShane 2021). Brain areas that play an important role in executive functions (memory, decision-making, attention, learning) such as the prefrontal cortex (PFC), striatum, and cerebellum, are sensitive to aging (Raz et al. 2003; Salat et al. 2004; Sullivan and Pfefferbaum 2006), as is the white matter connecting these areas (Raz et al. 2005).

Understanding changes in masticatory and swallowing performance in healthy aging is crucial for probing brain functions of aged individuals with cognitive impairment or dementia who also present with chewing and swallowing difficulties. Here, we study the effects of healthy aging on the tongue and jaw kinematics during natural feeding behavior in young and aged rhesus macaques. To do this, we precisely quantify 3*D* tongue and jaw movements using hi-resolution biplanar videoradiography.

## Methods

*Subjects:* Feeding experiments were performed on four male rhesus macaques (Macaca mulatta), of which three were young (age: *Y* = 7, *R* = 8, *B* = 11 years) and one old (*E* = 19 years), with each individual weighing between 9 −12 *kg*. All protocols were approved by the University of Chicago Animal Care and Use Committee and complied with the National Institutes of Health Guide for the Care and Use of Laboratory Animals.

*Behavioral task:* During the experiments, the subjects were seated on a primate chair with a restraint to avoid head movements during recording. The subjects were presented with food items, such as half grapes and gummy bears, attached to the end of a long stylus held by the experimenter. The food items differed in their liquid content and texture, with grapes being juicy while gummy bears being soft but dry and chewy. Each feeding trial lasted for a duration of 10 *s*, starting at food ingestion.

*Kinematics:* To record the tongue and jaw movements during natural feeding behavior, radio-opaque tantalum beads (1 *mm* diameter) were implanted in the tongue, mandible and the cranium while the subjects were under isoflurane anesthesia (2 −5%) (Laurence-Chasen et al. 2023). A superficial marker network of seven tantalum beads was placed just under the mucosal surface of the anterior two-thirds of the tongue, and a deep marker network of eight markers was placed deeper into the tongue (> 10 *mm*).

After the minimum recovery period of 2 weeks, the 3*D* position of jaw and tongue markers were recorded using high-resolution biplanar videoradiography (200 *Hz*). The data was then reconstructed using X-Ray Reconstruction of Moving Morphology (XROMM) workflow (Brainerd et al. 2010; Laurence-Chasen et al. 2020), which afforded a highly accurate and detailed analysis of the feeding behavior.

*Data analysis:* Feeding trials were divided and categorized based on gape cycle type, such as manipulation, stage 1 transport, left chew, right chew, stage 2 transport, intercalated swallow, and terminal swallow. This categorization was performed with the aid of X-ray videos, light camera videos, and XMALab’s biplanar marker tracking software (Knörlein et al. 2016). The 3*D* position of the anterior tongue and jaw markers were analyzed to determine differences in tongue and jaw movements between young and old subjects. Position data was calculated using an orthogonal coordinate system where the x-axis runs along the anteroposterior axis in the midline, the y-axis runs along the supero-inferior axis and the z-axis is oriented along the medio-lateral plane running through both the condyles (see (Olson et al. 2021) for more details). To remove the outliers from the data, we calculated the mean trajectory for each individual and removed the trajectories outside 5 standard deviations from the mean trajectory. Statistical analysis was performed to determine the significance of any observed differences in feeding behavior between the young and the old. The computational analysis was performed in MATLAB 2022*b* (Mathworks) and Python.

*Gape Cycles:* A gape cycle is defined as a complete jaw elevation-depression cycle measured in this study from maximum to maximum gape (Supplementary Fig. 1). A typical gape cycle is made up of four different gape cycle phases (Hiiemae et al. 1995). The gape cycle phases are determined by comparing the trajectory of superio-inferior position of the jaw (*J*_*y*_) with its second-derivative, jaw acceleration 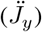. The gape cycle begins with the fast close phase (FC) when jaw acceleration hits maximum value, marking the start of maximum gape position. In this phase, the jaw is rapidly closed onto the food item. As the lower teeth meet either the food item or the upper teeth, jaw acceleration decreases to a minimum value. This is the beginning of the slow close or the power stroke (SC). During the SC phase, bite forces are applied on the food item. The SC phase ends when jaw position hits a maximum value, which marks the minimum gape. Towards the end of the SC phase, jaw acceleration also hits a minimum. After the minimum gape is achieved, the slow open phase (SO) begins, during which the jaw is slowly depressed to bring the tongue into contact with the food. SO phase ends and fast open (FO) phase begins as the mandible opening velocity increases again. FO ends as jaw acceleration hits the maximum value again, marking the end of maximum gape position.

## Results

### Tongue trajectories exhibit stereotypic patterns

The tongue plays a crucial role in positioning the bolus on the teeth row as the lower jaw moves up and down. As food is chewed and the bolus is formed, the tongue constantly adjusts its position within the oral cavity while maintaining its coordination with jaw motion (Palmer et al. 1997; Taniguchi et al. 2013). Tongue trajectories in younger subjects exhibited stereotypical movement patterns across hundreds of chew cycles (Fig. 1a-c). In contrast, they were considerably less stereotyped in older subjects (Fig. 1d). Moreover, the stereotypic tongue trajectories exhibited tight clustering. Therefore, to evaluate the degree of stereotypy of tongue trajectories, we calculated the frame-by-frame percentage of 3*D* tongue positions that fall within ±1 SD of the mean 3*D* position across all chew cycles. A high percentage represent tight clustering, and thus a greater stereotypy (Fig. 2a). We found that the clustering of tongue trajectories was modulated as a function of gape cycle. Tongue position showed tight clustering around minimum gape and both the start and end of maximum gape, as depicted by peak percentages in Fig. 2a. After the start of maximum gape, the trajectories initially spread out spatially then began to converge as the jaw approached minimum gape position. After minimum gape, the trajectories diverged again before clustering as the jaw reached the next maximum gape. This pattern was consistent across all subjects, suggesting a certain degree of stereotypy during chew cycles regardless of age. Peak values of tongue positions within ±1 SD of the mean 3*D* position ranged between 50 −70% in all young subjects and 40% for the aged subject at the minimum gape. Interestingly, there was a progressive decline in the clustering of tongue trajectories with increasing age of the animal, implying a progressive reduction in stereotypy (Fig. 2b, Pearson’s Correlation coefficient: 0.5126, *p* < 0.001).

**Figure 1.**
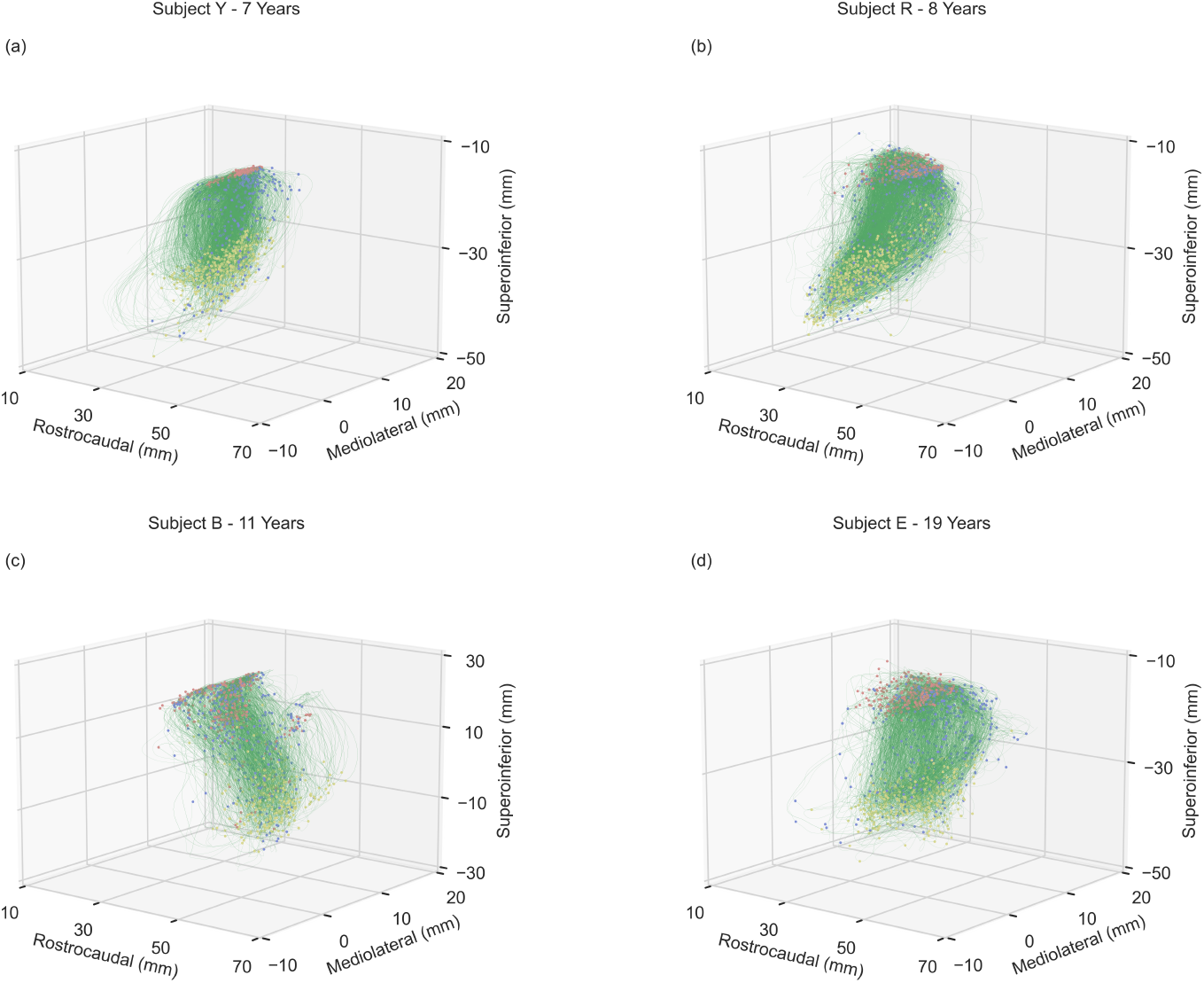
Stereotypic tongue motions during chews. 3*D* trajectories of the tongue tip are plotted ±0.5 *s* relative to minimum gape (red circles). Shown for younger subjects (a-c) and an older subject (d). Blue and yellow dots represent the maximum gape start and maximum gape end points, respectively, in each chewing cycle.

**Figure 2.**
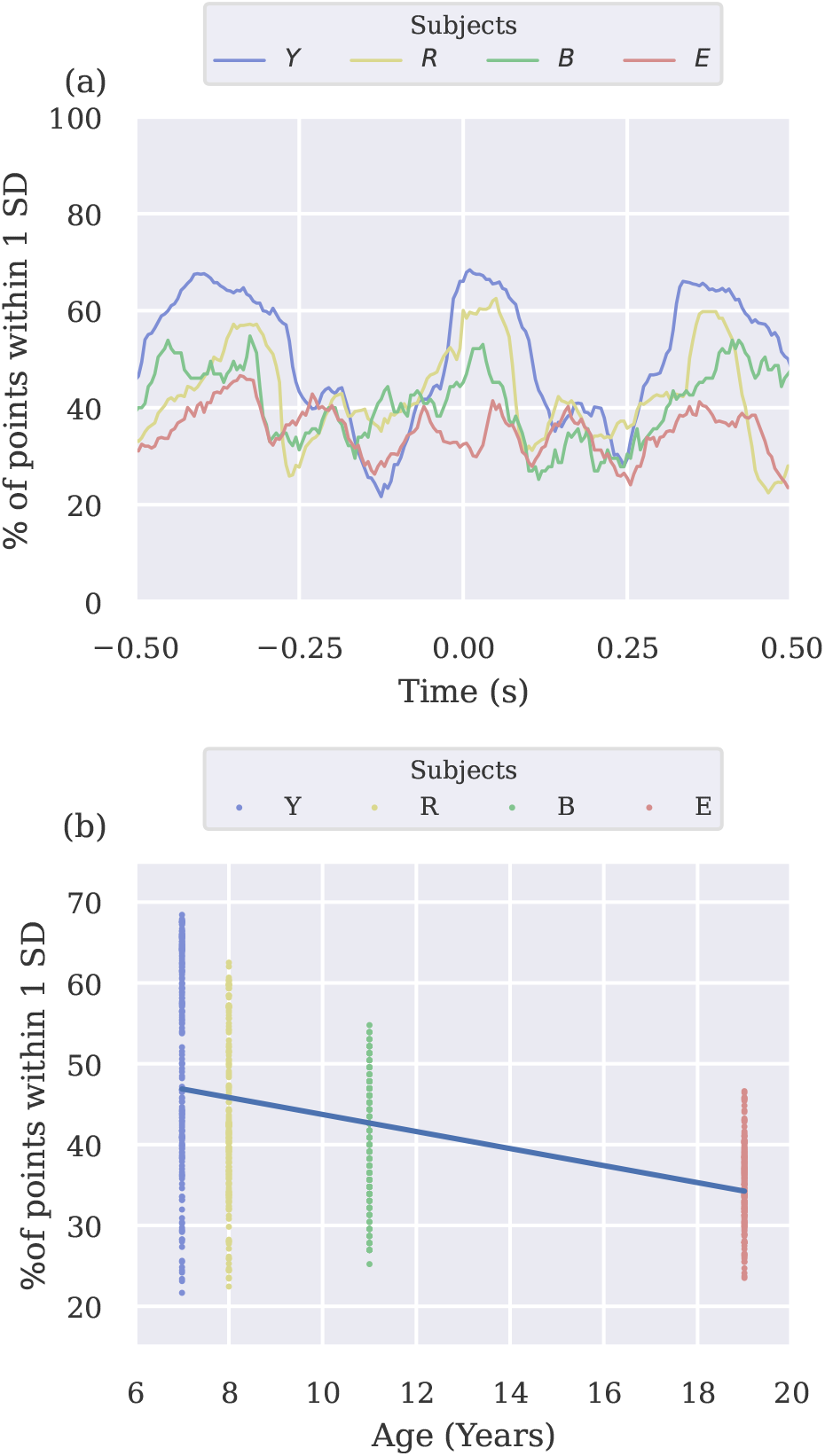
Clustering of 3D tongue trajectories during chew cycles. (a), Frame-by-frame percentage of 3*D* tongue positions within ±1 SD, shown for each subject as mean across chew cycles centered at minimum gape (0 *s*). (b), Percentage of 3*D* tongue positions within ±1 SD (data from Fig. 2a) plotted as a function of subject’s age. Solid line represents the linear fit.

### Characterization of tongue velocity during feeding

Next, we assessed the impact of aging on the velocity of tongue movements during chews as slowed movement has been associated with aging (Steele and Van Lieshout 2009). Tongue velocity exhibited significant modulation, i.e., difference between minimum and maximum velocities (Table 1, Fig. 3). Modulation and peak velocities were significantly higher in younger subjects than in the aged subject (Kruskal Wallis One Way ANOVA, *p* < 0.001, post-hoc paired comparisons: R vs E: *p* < 0.05, B vs E: *p* < 0.05). Peak velocities flanked the onset of minimum gape across all subjects. In younger subjects, tongue velocity reached first peak around 0.13 −0.15 *ms* prior to minimum gape, followed by deceleration as the jaw approached minimum gape, then by a fast acceleration to reach second peak velocity around 0.10 −0.15 *ms* after minimum gape. This temporal profile of modulation remained consistent across younger subjects. However, in the older subject, the peak velocities were more separated in time and minimum velocity occurred earlier.

**Table 1.** Mean and ±1 SD of minimum and maximum velocities of tongue tip across gape cycles and their corresponding mean modulation index.

| Subject | Minimum Velocity (cm/s) | Maximum Velocity (cm/s) | Modulation Index |
| --- | --- | --- | --- |
| Y | 0.0160 (0.0233) | 0.2189 (0.8444) | 0.2029 |
| R | 0.0703 (0.0658) | 0.2969 (0.1871) | 0.2266 |
| B | 0.0691 (0.0420) | 0.2775 (0.1654) | 0.2084 |
| E | 0.1140 (0.0563) | 0.2043 (0.0984) | 0.0903 |

**Figure 3.**
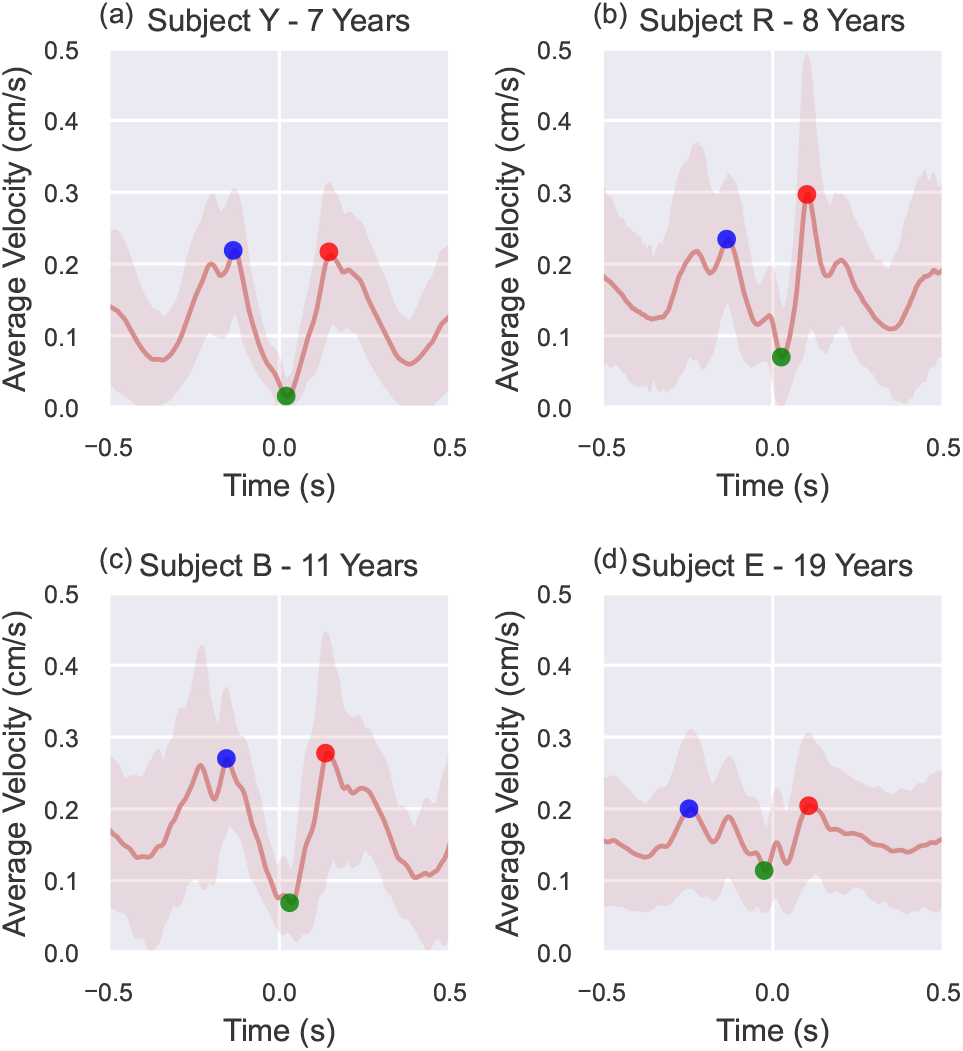
Average velocity of the tongue tip during chew cycles. The plots are centered ±0.5 *s* around minimum gape. Minimum average velocity is marked by green points and maximum average velocities before and after the minimum gape are marked by blue, and red points, respectively. Shaded regions denote ±1 SD. Shown for younger (a - c) and older (d) subjects.

### Coordination between tongue and jaw movements

To evaluate tongue-jaw coordination, we first examined the relationship between tongue and jaw positions during chew cycles. Figure 4a shows the rostro-caudal position of the tongue tip (i.e, protrusion-retrusion) and the supero-inferior position of the jaw (i.e, elevation-depression) as a function of percentage chew cycle. This normalization was necessary because of the variable duration of chew cycles within and across subjects. Different temporal profiles were found between young and aged subjects (Fig. 4a-d), suggesting a high degree of inter-subject variability. Nevertheless, across all subjects, peak tongue protrusion always lagged minimum gape. The mandible reached minimum gape around 50% of the chew cycle while the tongue reached its most protruded position around 60 −75% of the chew cycle. The temporal lag between peak position of the mandible and tongue was noticeably shorter in 2 out of 3 young subjects compared to the older subject (Kruskal Wallis One Way ANOVA, R vs E, B vs E, *p* < 0.05). Similar results were obtained when using mandible pitch (Supplementary Figure 4), but roll and yaw did not show considerable variations during chew cycles. In all subjects, there was a high correlation between tongue and jaw positions (Fig. 4e-h). However, the lag when peak correlations occurred differed between young and old subjects (Kruskal Wallis One Way ANOVA, *p* < 0.001); peak correlation was closer to zero lag (−35 to −25 *ms*, Fig. 4e-g) in young subjects and was at −90 *ms* lag in the aged subject (Fig. 4h).

**Figure 4.**
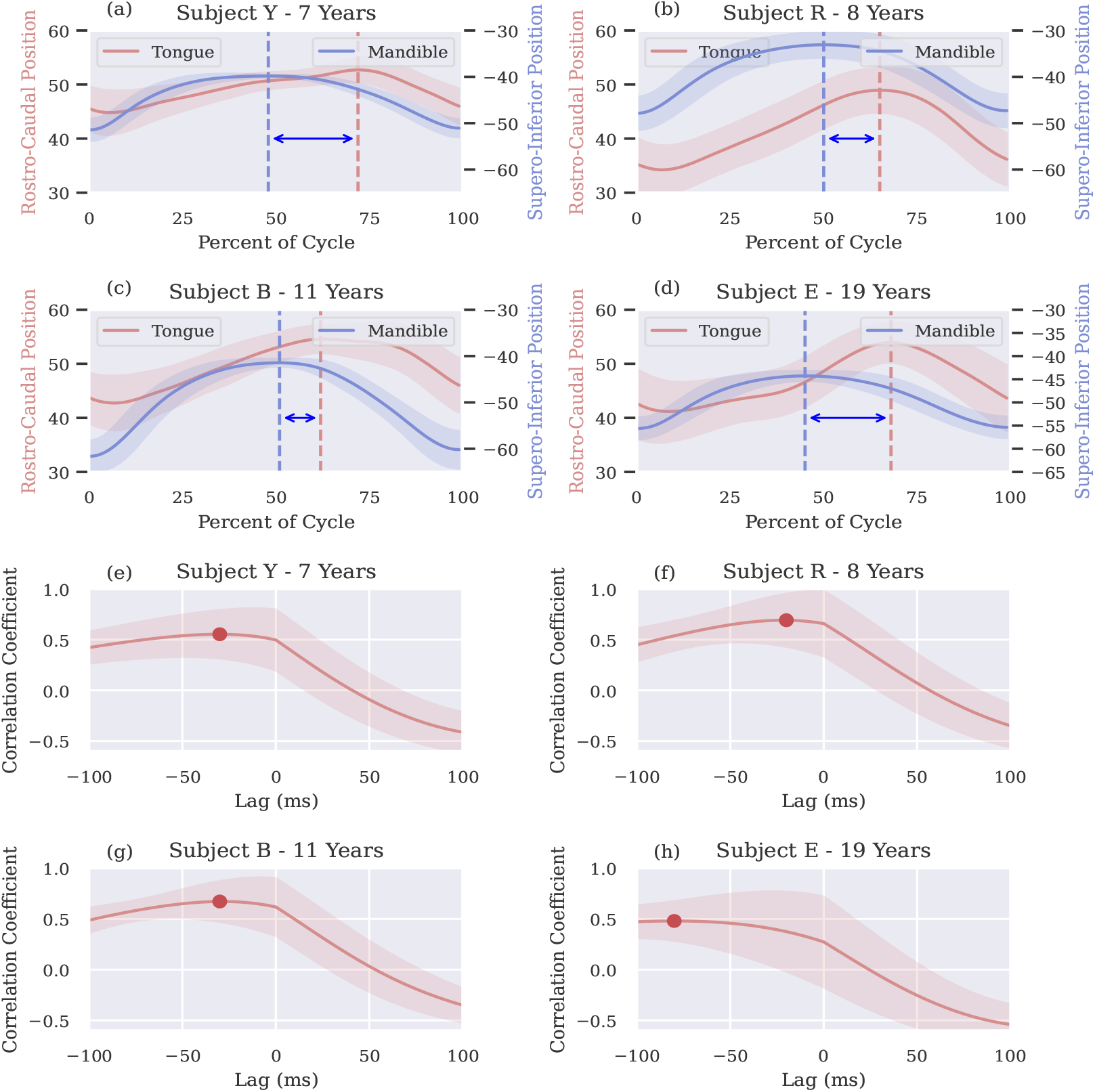
Relationship between tongue and jaw positions during chew cycles. (a-d), Mean tongue tip position in the rostro-caudal axis (left y-axis increases anteriorly) and mean mandible position in the supero-inferior axis (right y-axis increases superiorly) are plotted as a function of percentage chew cycle. The gape cycles start from maximum gape and the minimum gape around 50% of the cycle. The length of the double sided arrow indicates the lag between the time to peak tongue and mandible positions marked by dashed lines. (e-h), *Correlation between tongue (rostro-caudal) and jaw (supero-inferior) movements*. Plots show the cross-correlation coefficients at specific lags. Shown for each subject separately. Shaded regions denote ±1 SD.

### Characterization of gape cycle phases

We also evaluated the impact of aging on the frequency of gape cycles characterized as manipulation and/or stage I transport cycles, or cycles when swallows (both intercalated and terminal) occurred. Aging exerted a noticeable influence on the number of manipulation cycles and the swallow frequency; the older subject exhibited significantly higher average number of manipulation cycles compared to the two younger subjects when feeding on grapes (Fig. 5a), two-tailed t-test, R: *p* < 0.001, B: *p* < 0.01). While feeding on grapes, the older subject displayed increased variance in the average number of manipulation cycles compared to younger subjects (F-test, grape: Y and B: *p* < 0.01; gummy bear: R: *p* < 0.01, Y: *p* < 0.05). With swallows, the older subject exhibited a higher frequency compared to the younger subjects with gummy bear only (t-test, R vs E: *p* < 0.01, Y vs E: *p* < 0.001; F-test, *p* < 0.05). Lastly, the duration of chew cycles was higher in grape and more variable in both food types in the aged subject compared to most younger subjects (Fig. 5e, grape: R and Y, t-test, and F-test: *p* < 0.001; Fig. 5f, gummy bear: Y, t-test: *p* < 0.001, B, *p* < 0.01).

To understand how the gape cycle phases were affected by aging, we measured the normalized relative duration of the four gape cycle phases (Fig. 5g). The relative phase of the older subject differed significantly from the younger subjects. Specifically, the relative duration in the fast close and slow open increased significantly and the relative duration of the fast open reduced significantly in the older subject. We noticed that the relative increase in slow open was the largest in magnitude of all the changes in the older animal.

**Figure 5.**
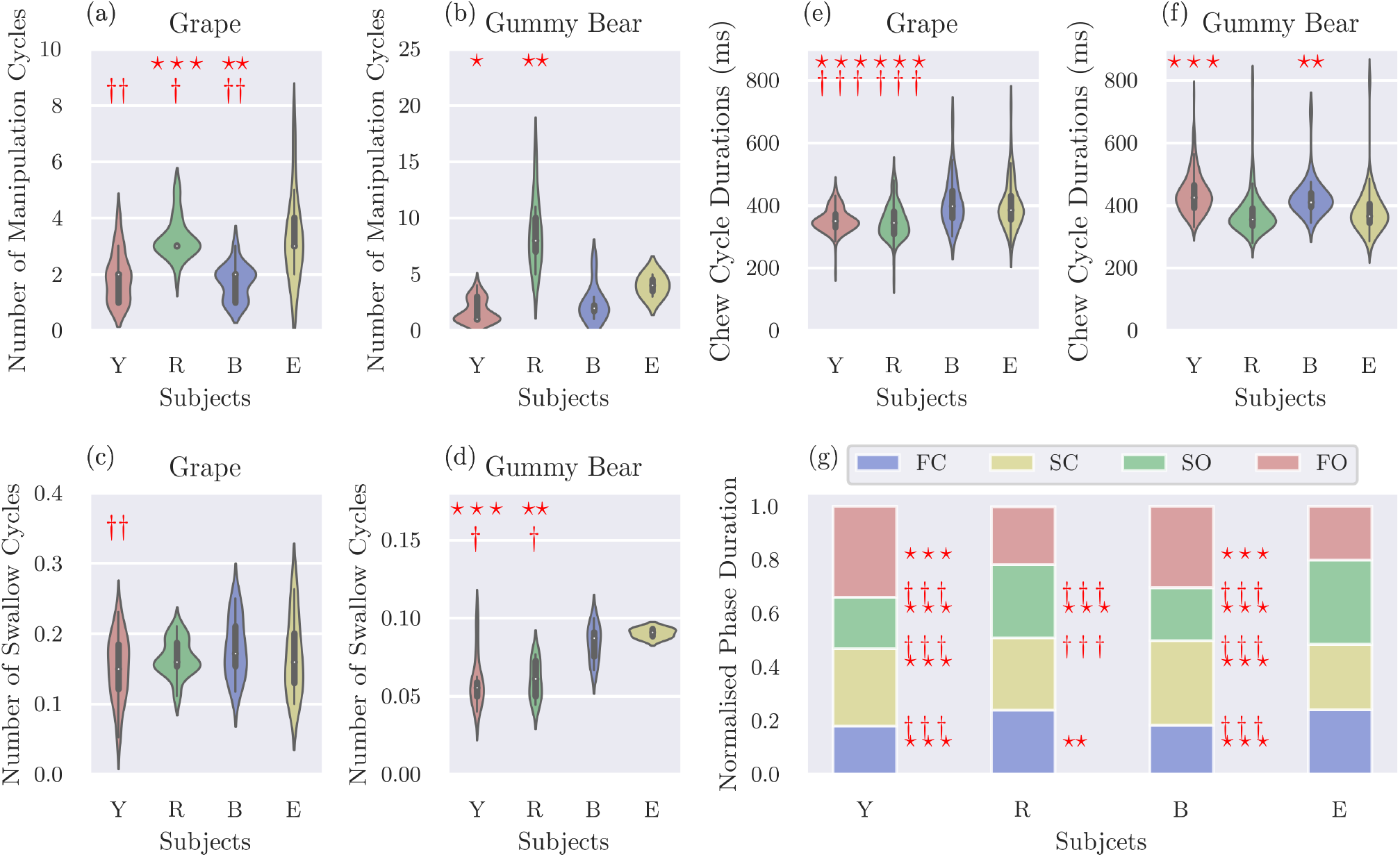
Frequencies of manipulation and swallow cycles shown per food type. (a-b), Number of manipulation and/or stage I transport cycles, prior to the onset of rhythmic chewing for grape and gummy bear. (c-d), Swallow frequency, as measured by number of swallows per 10 gape cycles for grape and gummy bear. (e-f), Chew cycle duration for grape and gummy bear, respectively. (g), *Relative duration of gape cycle phases*. FC is fast close, SC is slow close, SO is slow open, and FO is fast open phase. Since there were no significant food type effects, chews on both grapes and gummy bears were pooled for this analysis. Results of a two-tailed t-test and F-test of equality of variances (compared to the old subject) are indicated by asterisks and crosses, respectively: ⋆, † *p* < .05; ⋆⋆, †† *p* < .01; ⋆ ⋆ ⋆, † † † *p* < .001.

## Discussion

The aim of this study was to evaluate the impact of healthy aging on 3*D* tongue and jaw movements during natural feeding behavior in rhesus macaques. While feeding behavior was not severely impaired in our old subject, our analyses of tongue and jaw motion in 3*D* during feeding revealed important aspects in which feeding performance differed between our young and old subjects.

First, we found a substantial reduction in stereotypy of tongue trajectories as individuals age. Stereotypic movement patterns have been observed in arm reaching, walking, and skilled movements (Chiovetto and Giese 2013; Corbetta and Spencer 1996; Grillner and Zangger 1975a,b, 1979; Konczak’johannes et al. 1997; Marder and Calabrese 1996). It may be indicative of a mechanism to simplify the control of moving various joints as a synergy which requires a higher-level control of a movement pattern as opposed to controlling individual joints or muscles (Mussa-Ivaldi et al. 1985; Mussa-Ivaldi and Solla 2004). Younger individuals may demonstrate greater stereotypy when eating because of a more efficient motor control and coordination. Potential factors that may contribute to a reduced stereotypy are age-related changes in orofacial musculature, such as alterations in muscular strength, tone, endurance (Robbins et al. 1995; Welford 1984) and coordination (Steele and Van Lieshout 2009).

Our study also found a reduction in the modulation of tongue velocity with age, suggesting a lack of ability to fine-tune velocity. Past studies have found that fatigue and muscle weakness are associated with loss of muscle mass, speed, and strength found in aging (Cullins and Connor 2017; Welford 1984). Despite a reduction in tongue velocity with aging, the characteristic velocity curve patterns were preserved. This suggests that this may be an invariant aspect of chewing that is unaffected by aging.

Our research identified a high correlation between antero-posterior tongue and supero-inferior jaw movements in younger subjects, occurring close to zero lag. However, in older subject, the peak correlation shifted significantly away from zero lag, indicating reduced coordination between tongue and jaw movements. This deviation from zero lag is an important marker of healthy aging, highlighting age-related alterations in the coordination between tongue and jaw, crucial for functional activities such as speech and mastication (Masapollo et al. 2023; Thies et al. 2023).

Aging showed a food-dependent effect on swallow frequency, consistent with previous studies that reported age-associated decline in feeding performance Donini et al. (2003); Madhavan et al. (2016); Ney et al. (2009); Payne and Morley (2017, 2018). There were no significant changes in the swallow frequency during grape feeding. However, during gummy bear feeding, the older subject exhibited significant variance in swallow cycles compared to the younger subjects. It is interesting to note that the older subject had no difficulty in handling grapes with high liquid content whereas it had difficulty in dealing with chewy food type like the gummy bear. Diminished dental occlusion, tooth wear, or tooth loss may necessitate more extensive chewing to achieve adequate bolus breakdown (Peyron et al. 2018; Yven et al. 2006). Furthermore, alterations in tongue and jaw coordination, as well as reduced muscular efficiency, may contribute to the need for additional manipulation cycles to achieve optimal bolus consistency for swallowing. The difference in results observed for different food types illustrates the importance of including multiple food types in future studies.

The results of this study are to be interpreted with the following limitations in mind. Our study did not explore potential confounding factors such as individual variations in behavior, which could impact masticatory performance and coordination in aging individuals. Addressing this limitation in future research will provide a better understanding of the complexities of age-related changes in masticatory function. In conclusion, the knowledge gained from the study has important implications for the neuromechanical underpinnings of pathological aging such as in AD. Additionally, our findings may offer valuable groundwork for understanding earlier identification of individuals with chronic oral health issues who may be at risk for developing AD and other age related dementias (ARD) and for the development of effective interventions aimed at improving oral health outcomes in this group, with the potential to prevent the onset or the progression of AD/ARD.

## Author contributions

Conceptualization: F.I.A-M; Methodology: F.I.A-M; Software: S.P; Formal analysis: K.H, S.P., F.I.A-M; Investigation: F.I.A-M; Writing – Original draft: S.P.; Writing – Review & Editing: F.I.A-M, S.P.; Visualization: S.P; Supervision, Project Administration, and Funding Acquisition: F.I.A-M.

## Acknowledgements

We thank Dr. Jeffrey D. Laurence-Chasen for data collection and software, Victoria Hosack for XROMM data processing, Jared Luckas for initial data analysis and Dr. Jing-Sheng Li for valuable inputs and discussions.

## Supplemental material

**Supplementary Figure 1.**
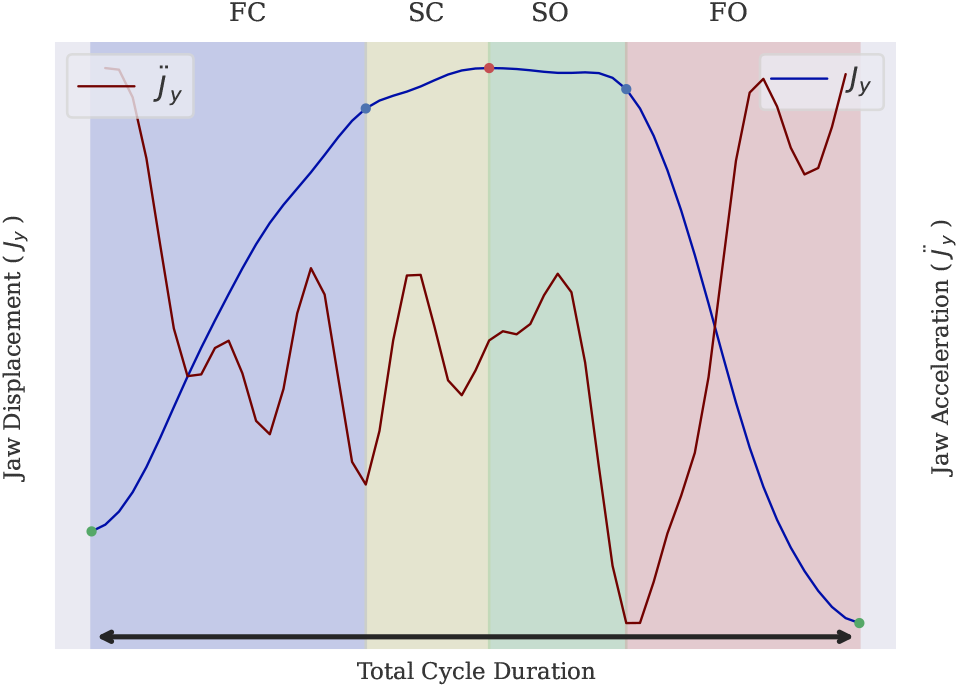
Illustration of the four phases of a gape cycle. Cycle duration is measured from maximum gape to the next maximum gape, shown as green circles on the jaw displacement (*J*_*y*_). The boundaries of the four chew phases, i.e., fast close (FC), slow close (SC), slow open (SO), and fast open (FO), are defined by jaw acceleration 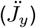 The transition from FC to SC (blue circle) is identified as the largest negative peak of 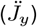 between the start of maximum gape and minimum gape (red circle). The second blue circle on *Jy* represents SO-FO transition based on the largest negative peak between minimum gape and maximum gape.

**Supplementary Figure 2.**
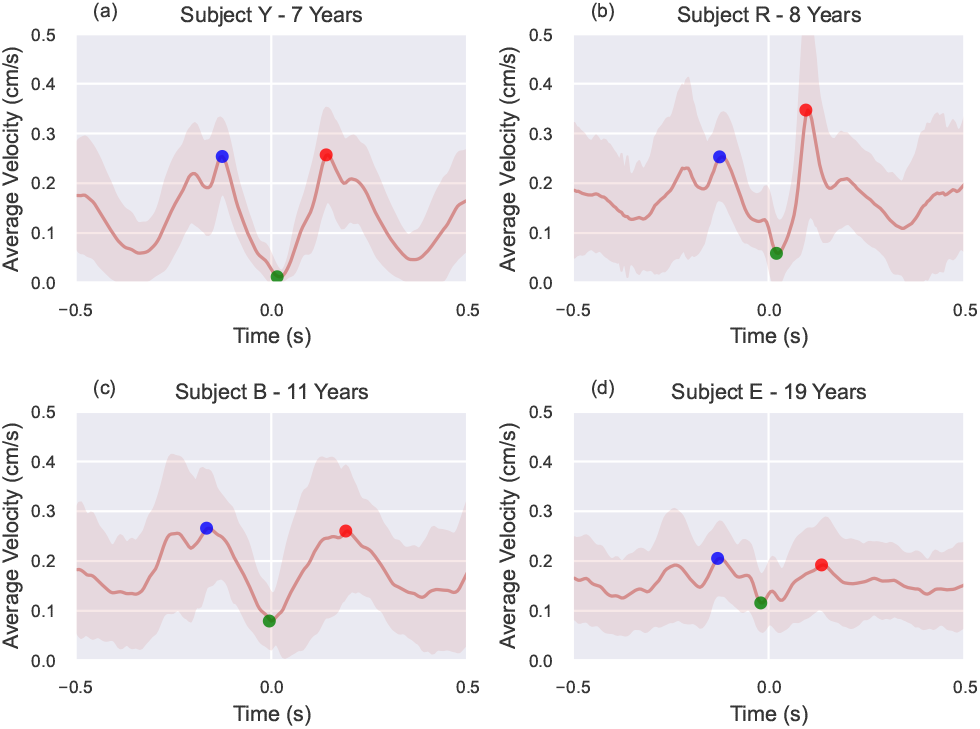
Average tongue velocity during grape chew cycles. The plots are centered ±0.5 *s* around the minimum gape. Minimum average velocities and maximum average velocities before and after the minimum gape are marked by green, blue, and red points, respectively. Shaded regions denote ±1 SD. Shown for younger (a - c) and old (d) subjects.

**Supplementary Figure 3.**
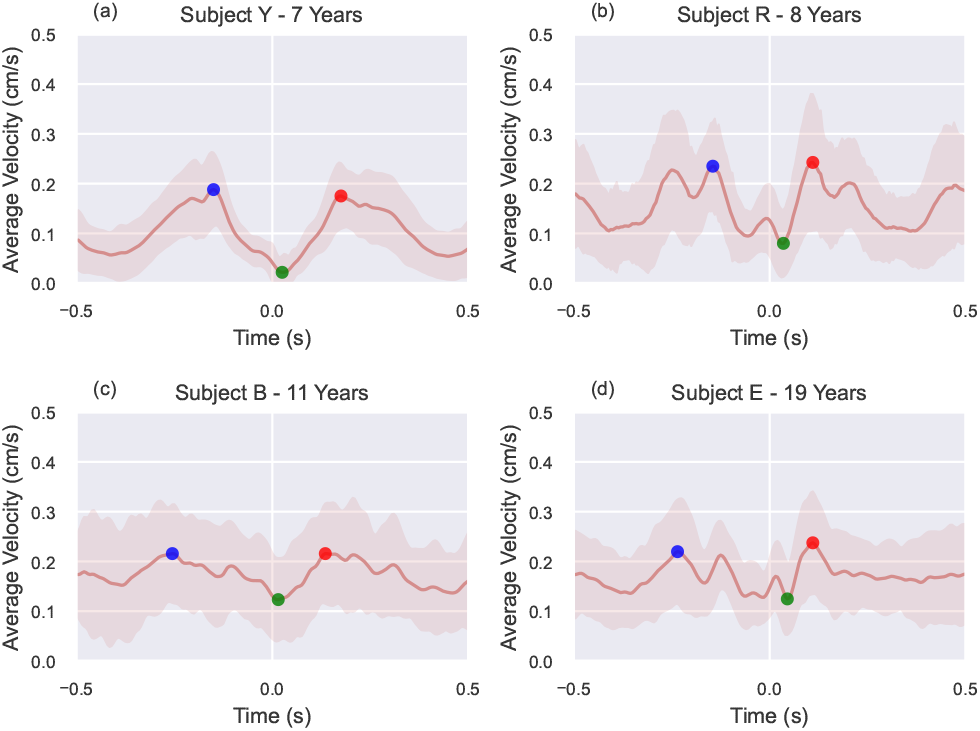
Average tongue velocity during gummy bear chew cycles. The plots are centered ±0.5 *s* around the minimum gape. Minimum average velocities and maximum average velocities before and after the minimum gape are marked by green, blue, and red points, respectively. Shaded regions denote ±1 SD. Shown for younger (a - c) and old (d) subjects.

**Supplementary Figure 4.**
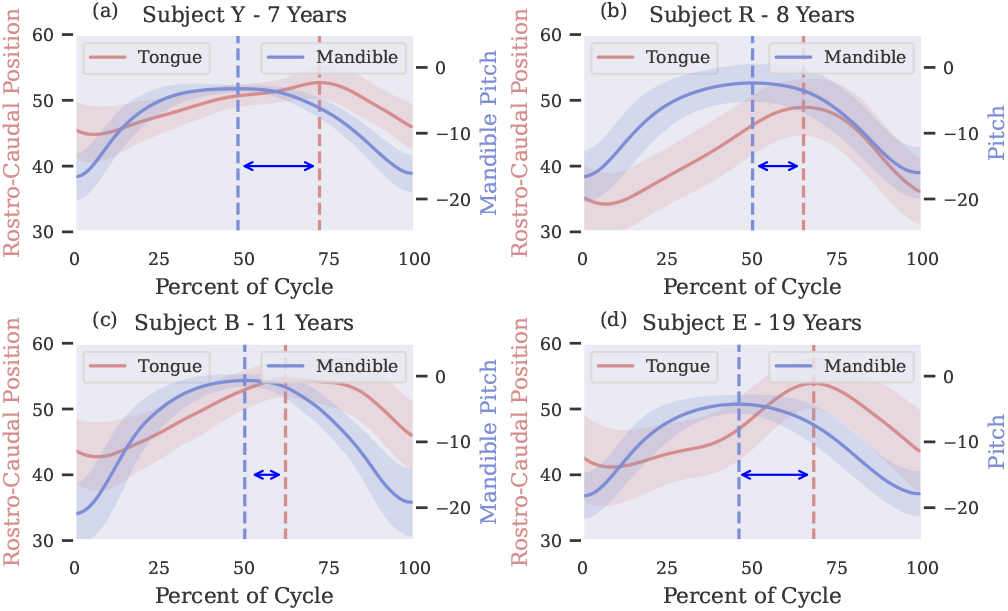
Relationship between tongue and jaw positions during chew cycles. Mean tongue tip position in the rostro-caudal axis (left y-axis increases anteriorly) and mean mandible pitch are plotted as a function of percentage chew cycle. The gape cycles start from maximum gape. Shaded regions denote ±1 SD. The length of the double sided arrow indicates the lag between the time to peak tongue and mandible positions marked by dashed lines.

## References

Arce-McShane FI. 2021. The association between age-related changes in oral neuromechanics and Alzheimer’s disease. Advances in geriatric medicine and research 3.

Avivi-Arber L, Sessle BJ. 2018. Jaw sensorimotor control in healthy adults and effects of ageing. Journal of oral rehabilitation 45:50–80.

Bakke M, Holm B, Jensen BL, Michler L, Møller E. 1990. Unilateral, isometric bite force in 8-68-year-old women and men related to occlusal factors. European Journal of Oral Sciences 98:149–158.

Bhandari A, Radhu N, Farzan F, Mulsant BH, Rajji TK, Daskalakis ZJ, Blumberger DM. 2016. A meta-analysis of the effects of aging on motor cortex neurophysiology assessed by transcranial magnetic stimulation. Clinical neurophysiology: official journal of the International Federation of Clinical Neurophysiology 127:2834–2845.

Brainerd EL, Baier DB, Gatesy SM, Hedrick TL, Metzger KA, Gilbert SL, Crisco JJ. 2010. X-ray reconstruction of moving morphology (XROMM): precision, accuracy and applications in comparative biomechanics research. Journal of Experimental Zoology Part A: Ecological Genetics and Physiology 313:262–279.

Brodoehl S, Klingner C, Stieglitz K, Witte OW. 2013. Age-related changes in the somatosensory processing of tactile stimulation– an fMRI study. Behavioural brain research 238:259–264.

Cheng CH, Chan PYS, Baillet S, Lin YY. 2015. Age-Related Reduced Somatosensory Gating Is Associated with Altered Alpha Frequency Desynchronization. Neural Plasticity 2015.

Cheng CH, Lin YY. 2013. Aging-related decline in somatosensory inhibition of the human cerebral cortex. Experimental brain research 226:145–152.

Chiovetto E, Giese MA. 2013. Kinematics of the Coordination of Pointing during Locomotion. PLOS ONE 8:e79555.

Corbetta D, Spencer JP. 1996. Development of Reaching during the First Year: Role of Movement Speed. Article in Journal of Experimental Psychology Human Perception & Performance.

Cullins MJ, Connor NP. 2017. Alterations of intrinsic tongue muscle properties with aging. Muscle & Nerve 56:E119–E125.

Daniels SK, Corey DM, Hadskey LD, Legendre C, Priestly DH, Rosenbek JC, Foundas AL. 2004. Mechanism of sequential swallowing during straw drinking in healthy young and older adults.

Dominy SS, Lynch C, Ermini F, Benedyk M, Marczyk A, Konradi A, Nguyen M, Haditsch U, Raha D, Griffin C, Holsinger LJ, Arastu-Kapur S, Kaba S, Lee A, Ryder MI, Potempa B, Mydel P, Hellvard A, Adamowicz K, Hasturk H, Walker GD, Reynolds EC, Faull RL, Curtis MA, Dragunow M, Potempa J. 2019. Porphyromonas gingivalis in Alzheimer’s disease brains: Evidence for disease causation and treatment with small-molecule inhibitors. Science Advances 5.

Donini LM, Savina C, di Roma U, Sapienza L, Roma IL, di Scienza dell l, Donini AM, Savina C, Can-mentazione C. 2003. Eating Habits and Appetite Control in the Elderly: The Anorexia of Aging. International Psychogeriatrics 15:73–87.

Fujiyama H, Hinder MR, Schmidt MW, Tandonnet C, Garry MI, Summers JJ. 2012. Age-related differences in corticomotor excitability and inhibitory processes during a visuomotor RT task. Journal of cognitive neuroscience 24:1253–1263.

Grillner S, Zangger P. 1975a. How detailed is the central pattern generation for locomotion? Brain Research 88:367.

Grillner S, Zangger P. 1975b. How detailed is the central pattern generation for locomotion? Brain Research 88:367.

Grillner S, Zangger P. 1979. On the central generation of locomotion in the low spinal cat. Experimental Brain Research 34:241–261.

Hatch JP, Shinkai RSA, Sakai S, Rugh JD, Paunovich ED. 2001. Determinants of masticatory performance in dentate adults. Archives of oral biology 46:641–648.

Heise KF, Zimerman M, Hoppe J, Gerloff C, Wegscheider K, Hummel FC. 2013. The aging motor system as a model for plastic changes of GABA-mediated intracortical inhibition and their behavioral relevance. The Journal of neuroscience: the official journal of the Society for Neuroscience 33:9039–9049.

Helkimo E, Carlsson GE, Helkimo M. 1977. Bite force and state of dentition. Acta odontologica scandinavica 35:297–303.

Heuninckx S, Wenderoth N, Swinnen SP. 2008. Systems neuroplasticity in the aging brain: recruiting additional neural resources for successful motor performance in elderly persons. The Journal of neuroscience: the official journal of the Society for Neuroscience 28:91–99.

Hiiemae K, Heath MR, Heath G, Kazazoglu E, Murray J, Sapper D, Hamblett K. 1996. Natural bites, food consistency and feeding behaviour in man. Archives of Oral Biology 41:175–189.

Hiiemae KM, Hayenga SM, Reese A. 1995. Patterns of tongue and jaw movement in a cinefluorographic study of feeding in the macaque. Archives of Oral Biology 40:229–246.

Hiiemae KM, Palmer J. 1999. Food transport and bolus formation during complete feeding sequences on foods of different initial consistency. Dysphagia 14:31–42.

Hutchinson S, Kobayashi M, Horkan CM, Pascual-Leone A, Alexander MP, Schlaug G. 2002. Age-related differences in movement representation. NeuroImage 17:1720–1728.

Kalisch T, Ragert P, Schwenkreis P, Dinse HR, Tegenthoff M. 2009. Impaired Tactile Acuity in Old Age Is Accompanied by Enlarged Hand Representations in Somatosensory Cortex. Cerebral Cortex 19:1530–1538.

Knörlein BJ, Baier DB, Gatesy SM, Laurence-Chasen JD, Brainerd EL. 2016. Validation of XMALab software for marker-based XROMM. Journal of Experimental Biology 219:3701–3711.

Kohyama K, Mioche L, Martin JF. 2002. Chewing patterns of various texture foods studied by electromyography in young and elderly populations. Journal of Texture Studies 33:269–283.

Konczakjohannes J, Dichgans KK, Konczak J Dichgans. 1997. The development toward stereotypic arm kinematics during reaching in the first 3 years of life. Exp Brain Res 117:346–354.

Koshino H, Hirai T, Ishijima T, Ikeda Y. 1997. Tongue motor skills and masticatory performance in adult dentates, elderly dentates, and complete denture wearers. The journal of prosthetic dentistry 77:147–152.

Laurence-Chasen JD, Manafzadeh AR, Hatsopoulos NG, Ross CF, Arce-McShane FI. 2020. Integrating XMALab and DeepLabCut for high-throughput XROMM. Journal of Experimental Biology 223:jeb226720.

Laurence-Chasen JD, Ross CF, Arce-McShane FI, Hatsopoulos NG. 2023. Robust cortical encoding of 3D tongue shape during feeding in macaques. Nature Communications 2023 14:1 14:1–10.

Lenz M, Tegenthoff M, Kohlhaas K, Stude P, Höffken O, Tossi MA, Kalisch T, Dinse HR. 2012. Increased excitability of somatosensory cortex in aged humans is associated with impaired tactile acuity. The Journal of neuroscience: the official journal of the Society for Neuroscience 32:1811–1816.

Logemann JA. 1990. Effects of aging on the swallowing mechanism. Otolaryngologic Clinics of North America 23:1045–1056.

Lund JP, Kolta A. 2006. Generation of the central masticatory pattern and its modification by sensory feedback. Dysphagia 21:167–174.

Madhavan A, Lagorio LA, Crary MA, Dahl WJ, Carnaby GD. 2016. Prevalence of and risk factors for dysphagia in the community dwelling elderly: A systematic review. Journal of Nutrition, Health and Aging 20:806–815.

Marder E, Calabrese RL. 1996. Principles of rhythmic motor pattern generation. 10.1152/physrev.1996.76.3.687 76:687–717.

Masapollo M, Nittrouer S, Stepp Editor CE, Kent RD. 2023. Interarticulator Speech Coordination: Timing Is of the Essence. Journal of Speech, Language, and Hearing Research 66:901–915.

McGinley M, Hoffman RL, Russ DW, Thomas JS, Clark BC. 2010. Older adults exhibit more intracortical inhibition and less intracortical facilitation than young adults. Experimental gerontology 45:671–678.

Mistry S, Hamdy S. 2008. Neural control of feeding and swallowing. Physical medicine and rehabilitation clinics of North America 19:709–728.

Morquette P, Lavoie R, Fhima MD, Lamoureux X, Verdier D, Kolta A. 2012. Generation of the masticatory central pattern and its modulation by sensory feedback. Progress in neurobiology 96:340–355.

Mussa-Ivaldi FA, Hogan N, Blzzl E. 1985. Neural, Mechanical, and Geometric Factors Subserving Arm Posture in Humans’. The Journal of Neuroscience 5:2732–2743.

Mussa-Ivaldi FA, Solla SA. 2004. Neural primitives for motion control. IEEE Journal of Oceanic Engineering 29:640–650.

Nakamura Y, Katakura N. 1995. Generation of masticatory rhythm in the brainstem. Neuroscience research 23:1–19.

Newton JP. 1987. Changes in human masseter and medial pterygoid muscles with age: a study by computed tomography. Gerodontology 3:151–154.

Newton JP, Yemm R, Abel RW, Menhinick S. 1993. Changes in human jaw muscles with age and dental state. Gerodontology 10:16–22.

Ney DM, Weiss JM, Kind AJ, Robbins J. 2009. Senescent swallowing: Impact, strategies, and interventions. Nutrition in Clinical Practice 24:395–413.

Olson RA, Montuelle SJ, Chadwell BA, Curtis H, Williams SH. 2021. Jaw kinematics and tongue protraction–retraction during chewing and drinking in the pig. Journal of Experimental Biology 224:jeb239509.

Österberg T, Carlsson GE, Tsuga K, Sundh V, Steen B. 1996. Associations between self-assessed masticatory ability and some general health factors in a Swedish population. Gerodontology 13:110–117.

Paganini-Hill A, White SC, Atchison KA. 2012. Dentition, Dental Health Habits, and Dementia: The Leisure World Cohort Study. Journal of the American Geriatrics Society 60:1556–1563.

Palmer JB, Hiiemae KM, Liu J. 1997. Tongue-jaw linkages in human feeding: a preliminary videofluorographic study. Archives of Oral Biology 42:429–441.

Payne M, Morley JE. 2018. Dysphagia, Dementia and Frailty. Journal of Nutrition, Health and Aging 22:562–565.

Payne MA, Morley JE. 2017. Dysphagia: A New Geriatric Syndrome. Journal of the American Medical Directors Association 18:555–557.

Peyron MA, Blanc O, Lund JP, Woda A. 2004. Influence of age on adaptability of human mastication. Journal of neurophysiology 92:773–779.

Peyron MA, Santé-Lhoutellier V, François O, Hennequin M. 2018. Oral declines and mastication deficiencies cause alteration of food bolus properties. Food & Function 9:1112–1122.

Prinz JF, Lucas PW. 1997. An optimization model for mastication and swallowing in mammals. Proceedings of the Royal Society of London. Series B: Biological Sciences 264:1715–1721.

Raz N, Lindenberger U, Rodrigue KM, Kennedy KM, Head D, Williamson A, Dahle C, Gerstorf D, Acker JD. 2005. Regional brain changes in aging healthy adults: general trends, individual differences and modifiers. Cerebral cortex 15:1676–1689.

Raz N, Rodrigue KM, Kennedy KM, Head D, Gunning-Dixon F, Acker JD. 2003. Differential aging of the human striatum: longitudinal evidence. American Journal of Neuroradiology 24:1849–1856.

Robbins JA, Levine R, Wood J, Roecker EB, Luschei E. 1995. Age Effects on Lingual Pressure Generation as a Risk Factor for Dysphagia. Journal of Gerontology: Series A 50:257–262.

Rogus-Pulia N, Malandraki GA, Johnson S, Robbins JA. 2015. Understanding Dysphagia in Dementia: The Present and the Future. Current Physical Medicine and Rehabilitation Reports 3:86–97.

Rueda-Delgado LM, Heise KF, Daffertshofer A, Mantini D, Swinnen SP. 2019. Age-related differences in neural spectral power during motor learning. Neurobiology of aging 77:44–57.

Salat DH, Buckner RL, Snyder AZ, Greve DN, Desikan RSR, Busa E, Morris JC, Dale AM, Fischl B. 2004. Thinning of the cerebral cortex in aging. Cerebral cortex 14:721–730.

Schindler HJ, Stengel E, Spiess WEL. 1998. Feedback control during mastication of solid food textures—a clinical-experimental study. The Journal of prosthetic dentistry 80:330–336.

Sebastián M, Ballesteros S. 2012. Effects of normal aging on event-related potentials and oscillatory brain activity during a haptic repetition priming task. NeuroImage 60:7–20.

Secişl Y, Arici İncesu TK, Gürgör N, Beckmann Y, Ertekin C. 2016. Dysphagia in Alzheimer’s disease. Neurophysiologie Clinique/Clinical Neurophysiology 46:171–178.

Seidler R, Erdeniz B, Koppelmans V, Hirsiger S, Mérillat S, Jäncke L. 2015. Associations between age, motor function, and resting state sensorimotor network connectivity in healthy older adults. NeuroImage 108:47–59.

Sessle BJ. 2006. Mechanisms of oral somatosensory and motor functions and their clinical correlates. Journal of oral rehabilitation 33:243–261.

Steele CM, Van Lieshout P. 2009. Tongue Movements During Water Swallowing in Healthy Young and Older Adults. Journal of Speech, Language, and Hearing Research 52:1255–1267.

Sullivan EV, Pfefferbaum A. 2006. Diffusion tensor imaging and aging. Neuroscience & Biobehavioral Reviews 30:749–761.

Talelli P, Ewas A, Waddingham W, Rothwell JC, Ward NS. 2008. Neural correlates of age-related changes in cortical neurophysiology. NeuroImage 40:1772–1781.

Taniguchi H, Matsuo K, Okazaki H, Yoda M, Inokuchi H, Gonzalez-Fernandez M, Inoue M, Palmer JB. 2013. Fluoroscopic evaluation of tongue and jaw movements during mastication in healthy humans. Dysphagia 28:419–427.

Thies T, Mücke D, Ding A, Mefferd A. 2023. Phase relations between the tongue body and the jaw across rate modifications in younger and older speakers. Speech Production and Speech Physiology.

Ueno M, Yanagisawa T, Shinada K, Ohara S, Kawaguchi Y. 2008. Masticatory ability and functional tooth units in Japanese adults. Journal of oral rehabilitation 35:337–344.

Ward NS, Frackowiak RS. 2003. Age-related changes in the neural correlates of motor performance. Brain: a journal of neurology 126:873–888.

Watanabe Y, Hirano H, Matsushita K. 2015. How masticatory function and periodontal disease relate to senile dementia. Japanese Dental Science Review 51:34–40.

Wayler AH, Chauncey HH. 1983. Impact of complete dentures and impaired natural dentition on masticatory performance and food choice in healthy aging men. The Journal of prosthetic dentistry 49:427–433.

Welford AT. 1984. Between bodily changes and performance: Some possible reasons for slowing with age. Experimental Aging Research 10:73–88.

Yoshikawa M, Yoshida M, Nagasaki T, Tanimoto K, Tsuga K, Akagawa Y, Komatsu T. 2005. Aspects of swallowing in healthy dentate elderly persons older than 80 years. The Journals of Gerontology Series A: Biological Sciences and Medical Sciences 60:506–509.

Yven C, Bonnet L, Cormier D, Monier S, Mioche L. 2006. Impaired mastication modifies the dynamics of bolus formation. European journal of oral sciences 114:184–190.

